# Reversible Loss of Hippocampal Function in a Mouse Model of Demyelination/Remyelination

**DOI:** 10.1101/866723

**Authors:** Aniruddha Das, Chinthasagar Bastian, Lexie Trestan, Jason Suh, Tanujit Dey, Bruce Trapp, Selva Baltan, Hod Dana

**Author notes:** **Correspondence:** Hod Dana.

## Abstract

Demyelination of axons in the central nervous system (CNS) is a hallmark of multiple sclerosis (MS) and other demyelinating diseases. Cycles of demyelination, followed by remyelination, appear in the majority of MS patients, and are associated with the onset and quiescence of disease-related symptoms, respectively. Previous studies have shown in human patients and animal models that vast demyelination is accompanied by wide-scale changes to brain activity, but details of this process are poorly understood. We use electrophysiological recordings and nonlinear imaging of fluorescence from genetically-encoded calcium indicators to monitor the activity of hippocampal neurons during demyelination and remyelination processes over a period of 100 days. We find *in vitro* that synaptic transmission in CA1 neurons is diminished, and *in vivo* both CA1 and dentate gyrus (DG) neuronal firing rates are substantially reduced during demyelination and partially recover after a short remyelination period. This new approach allows monitoring how synaptic transmission changes, induced by cuprizone diet, are affecting neuronal activity, and can potentially be used to study the effects of therapeutic interventions in protecting the functionality of CNS neurons.

## Introduction

Multiple sclerosis (MS) is an inflammatory demyelinating disease of the central nervous system (CNS) that affects more than 2.3 million patients worldwide [1]. For approximately 85% of patients, disease progression begins with a “relapsing-remitting” phase, where symptoms including motor deficits and vision loss correlate with increased formation of demyelinated lesions detected by magnetic resonance imaging (MRI). Over time, these symptoms resolve into periods of clinical quiescence and partial remyelination of demyelinated axons [2; 3]. In the majority of MS patients, the disease course eventually transforms from the relapsing-remitting phase to a secondary-progressive phase, where symptoms continually worsen [4]. Traditionally, MS was considered to be mainly a white-matter disease, but today it is widely accepted that there is also a critical gray-matter component, including demyelination of the cerebral cortex and deep gray matter [5]. Previous studies have also identified changes in brain activity patterns of MS patients during both the relapsing-remitting and secondary-progressive forms of the disease, where presumably the reduced activity of MS-affected brain regions is partially compensated for by increased activity in other brain regions [6; 7].

Studying the relationship between demyelination/remyelination cycles, motor and cognitive deficits, and changes in brain activity patterns requires the use of appropriate animal models and proper experimental methods. A commonly-used mouse model for studying demyelination and remyelination is the cuprizone model [8; 9], in which the copper chelator cuprizone is added to the mouse diet and causes damage to oligodendrocyte cells in the CNS. The result is extensive demyelination in various brain areas, including the hippocampus, corpus callosum, cortex, and cerebral white matter, which is partially reversible upon cessation of the cuprizone diet [8; 10; 11; 12]. Interestingly, cuprizone-treated mice, similar to MS patients, were identified as having demyelination-induced reorganization of brain circuitry [13]. Moreover, the extensive brain demyelination resulted in significantly poorer performance in various behavioral tests, including hippocampus-dependent tasks [14]. This reduction in performance was reversed upon remyelination [15]. Therefore, the cuprizone model seems to capture the demyelination-induced brain-wide activity reorganization that has been identified in patients. However, the brain mapping methods used for identifying these brain-wide activity changes in living mice were usually functional MRI and EEG, which have limited spatial and temporal resolutions that restrict their ability to explore rapid and local changes in brain activity. In order to uncover demyelination-induced effects on single cells and local networks, a method which allows micron-scale identification and single action potential (AP) detection *in vivo* is required.

Substantial improvements in spatiotemporal resolution can be achieved by using nonlinear optics to detect brain activity [16]. Two-photon laser scanning microscopy (TPLSM) of the fluorescence signal from genetically-encoded calcium indicators (GECIs) enables monitoring the activity of thousands of individual neurons in the rodent brain with micron-scale resolution and high signal-to-noise ratio [17; 18; 19]. Moreover, state-of-the-art GECIs provide the temporal dynamics and sensitivity that are necessary to allow detection of single APs over time scales of months, mainly by switching to stable transgenic expression strategies [18; 19; 20; 21; 22; 23]. Therefore, this approach may enable the exploration of changes in brain activity following demyelination and remyelination with the required sensitivity and accuracy to detect changes in individual neurons over time.

In this study, we combined electrophysiology and TPLSM recording of acute and chronic activity patterns of neurons in the mouse hippocampus before, during, and after the mice were fed with cuprizone diet. We identified rapid degeneration of CA1 and DG activity following the initiation of cuprizone diet, which were faster than expected by the induced demyelination alone, which decreased neuronal activity to ~20% of its pre-cuprizone level. We also found that the synaptic transmission in CA1 neurons in response to sustained pre-synaptic stimuli was drastically diminished after the cuprizone diet initiation. Synaptic transmission and neuronal activity were partially recovered upon cessation of the cuprizone diet. Thus, our study presents a new experimental approach for studying the effects of toxin-induced changes to the brain physiology, such as demyelination and remyelination, on the integrity of individual neurons and neuronal circuits, and provides a platform to test the benefits of potential treatment options.

## Materials and Methods

All surgical and experimental procedures were conducted in accordance with protocols approved by the Lerner Research Institute Institutional Animal Care and Use Committee and Institutional Biosafety Committee.

### Hippocampal slice electrophysiology

Electrophysiological experiments were performed in the CA1 region of 400 µm-thick transverse hippocampal slices. A cohort of 46 mice were divided to control (n=18, normal diet, daily Rapamycin injection) and experimental groups (n=28, cuprizone diet with daily Rapamycin injections). Among those in experimental group, synaptic transmission was monitored and quantified beginning at 1 (n=2), 3 (n=9), 4 (n=8), 5 (n=3) and 6 weeks (n=3) of demyelination. The last group consisted of mice on cuprizone diet for 6 weeks which then were allowed 6 weeks on control diet to remyelinate (n=3, 6 W DM + 6 W RM). Synaptic transmission was recorded in Control (n=8, 0 week) and at 4 (n=5, 4 W C) and 6 weeks (n= 5, 6 W C) to match respective demyelination groups. Mice were decapitated after anesthesia and brains were immediately removed and placed in ice-cold (±4°C) artificial cerebrospinal fluid (ACSF). ACSF contained in mM: 126 NaCl, 3.5 KCl, 1.3 MgCl_2_, 2 CaCl_2_, 1.3 NaH_2_PO_4_, 25 NaHCO_3_, and 10 glucose at pH 7.4. Hippocampi were quickly dissected and sliced with a McIlwain tissue chopper. Transverse slices were then transferred to ACSF solution and were continuously bubbled with 95% O_2_/5% CO_2_ for at least 1-2h at room temperature to stabilize.

Individual slices were then transferred and placed on a Haas-type tissue chamber (Warner Instruments, Harvard Apparatus) on a custom mesh (Warner Instruments, Catalog # 99-1129) to avoid air bubble accumulation and were stabilized using a slice holder (Warner Instruments, Cat # 64-1415). Slices were kept at the interface of a moist humidified gas mixture of 95% O_2_/5% CO_2_ and warm ACSF (at 33-34°C) bubbled with 95% O_2_/5% CO_2_ continually at a flow rate of 3-3.5 ml/min. For simultaneous extracellular recordings of afferent volleys (AV) and excitatory post-synaptic potentials (EPSPs), glass microelectrodes with 1-micron tips were pulled (Sutter Instruments, Borosilicate Glass with filament) and filled with 2M NaCl (resistance 1-2 MΩ). Electrode tips were polished using a Micro-Forge (Leica, MF-830, Narishige, Japan) and placed in the CA1 region of stratum radiatum layer to record synaptic transmission. Responses were evoked by stimulation of Schaffer collateral commissural fibers by a custom-designed bipolar tungsten wire electrode (Microprobes Part# PI2ST30.1B3) with 50 µs-long pulses at intervals of 30 seconds.. After stabilization, synaptic transmission was recorded for at least 30 min as baseline. The changes in synaptic transmission properties were quantified by measuring the peak amplitude of the EPSPs, as well as their area (Clampfit 10.7) to account for smaller slower responses. Maximum EPSPs, evoked by 100% of the stimulation amplitude (1 mA), were chosen for these measurements.

### Installation of hippocampal windows

For hippocampal window implantation, we slightly modified a previously-described procedure [24; 25]. Thy1-jRGECO1a line 8.31 mice were anesthetized using isoflurane (2.5-3% for induction, 1.5% during surgery) on a 37°C heating pad. Local pain medication (bupivacaine 0.5%) was injected into the craniotomy area before the skin and connective tissue were removed to expose the skull. A circular craniotomy (3.2 mm diameter) was drilled using a hand drill (OMNIDRILL35, World Precision Instruments). The cortex and corpus callosum, but not the alveus above the dorsal part of the hippocampus, were removed by gentle suction with saline using 26G and 28G sharp and blunt needles connected to a vacuum line. After the hippocampus was exposed, it was kept in place by gently pressing a glass coverslip to it (3 mm diameter #1 glass, Warner Instruments) attached to a metal canula (2.65mm internal diameter, 3.2mm outer diameter, 1.8mm length, New England Small Tube Corp., type 304L stainless steel). The cannula was attached to the skull bone using dental cement (Contemporary Ortho-Jet, Lang Dental). A custom stainless-steel head post was cemented to the skull using the same dental cement.

### Recording of hippocampal activity

Seven to fourteen days after the craniotomy surgery, we started monitoring hippocampal activity. The same procedure was followed for all mice before, during, and after the cuprizone diet period. Mice were anesthetized (3% isoflurane) and injected with Chlorprothixene Hydrochloride (IM, 30 μl of a 0.33 mg/ml solution, Santa Cruz). Recordings started at least 30 min after injection, and isoflurane levels were reduced to 0.5-0.75% prior to recording to bring the mice to a lightly-anesthetized state, where they do not move but are sensitive to pain. Mice were put on a 37°C heating pad in the dark, and spontaneous activity was recorded using 20-120 mW of 1100nm excitation light (Insight X3, Spectra-Physics) and a resonant-scanner two-photon microscope (512×512 pixels, 30Hz acquisition rate, Bergamo II, Thorlabs). Light intensity was controlled using a Pockels Cell (model 350-105, Conoptics) to maintain similar signal-to-noise ratios across different animals. Data were recorded on a weekly basis from all animals, and each recording session included acquisition of 200 s for each field of view (FOV), 3-5 FOVs from CA1 and 2-4 FOVs from DG for each mouse.

### Data analysis

Data analysis was performed using custom Matlab scripts (Mathworks). Every two frames in the recorded time series were averaged to improve the signal-to-noise ratio, reducing the acquisition rate to 15Hz. Small drifts and movements of the imaged area during the 200 s recording time were corrected with a Matlab script using the TurboReg plug-in of ImageJ [26]. Ring-shaped regions of interest (ROIs) corresponding to all identifiable somata were selected using a semi-automatic graphical user interface [17]. Fluorescence signal from all pixels inside of a ROI were averaged to calculate the fluorescence signal (F) for every cell. Baseline fluorescence levels (F_0_) for each ROI were estimated as its F median value across the 200 s of recording. To estimate the AP firing from the fluorescence data, we calculated ΔF/F_0_=(F-F_0_)/F_0_ for each cell, and fed it to a published model [27] for extracting AP activity from calcium traces (for accepting the model assumption, a trace of 1+ΔF/F_0_ was used). We fit the model with the following parameters, according to jRGECO1a published characterization [19]: 1AP amplitude of 15%, decay time of 175ms, and Hill coefficient of 1.9. Saturation level was set to 0.001 and signal drift to 0.03. Noise level was estimated using the model’s internal function. The model results were inspected visually for accuracy, and for CA1 data the noise level estimation was increased by 50% to lower the false-positive rate. We identified active cells as cells with at least one identified AP by the AP-extraction model. The average activity was calculated as the total number of APs divided by the recording duration of 200s. An activity burst was defined as firing of at least 5 APs within 660 ms.

For calculating the cellular contrast, we averaged all pixels in the ring-shaped ROI (covering the cellular cytosol), and averaged 5 x 5 grid of pixels around the ROI center, corresponding to an area inside the nucleus. The contrast was calculated as the difference between the averaged cytosolic and nucleic signals, divided by the nucleic signal.

### Statistical analysis

Statistical analysis was performed using Matlab and R software (version 3.5.1; https://cran.r-project.org). We performed four different statistical tests to identify changes in activity across different recording dates and mice. Wilcoxon Rank Sum test was used to compare changes in median firing rate between two different recording dates from the same mice. Mann-Kendall Trend Test was used to identify if there is a monotonic trend in the median firing rates during the cuprizone diet period (8 recording dates) for individual mice. Paired t-test was used to compare change of median firing rates of cuprizone or control groups across two recording dates. A repeated measures linear mixed effects model (repeated ANOVA test) was performed to determine the interaction of group (cuprizone and control) and time (8 recording dates) on CA1 and DG median activity rates, separately. All p-values are reported in the manuscript, the level of significance was set to 5% for all hypothesis testing procedures.

For analysis of *in vitro* hippocampal experiments, all means are given ±S.E.M. and the significance of differences within a group was assessed by One-way ANOVA, followed by Bonferroni’s test. N denotes animal numbers in the text and parenthesis in figure 1 indicates slice numbers.

**Figure 1.**
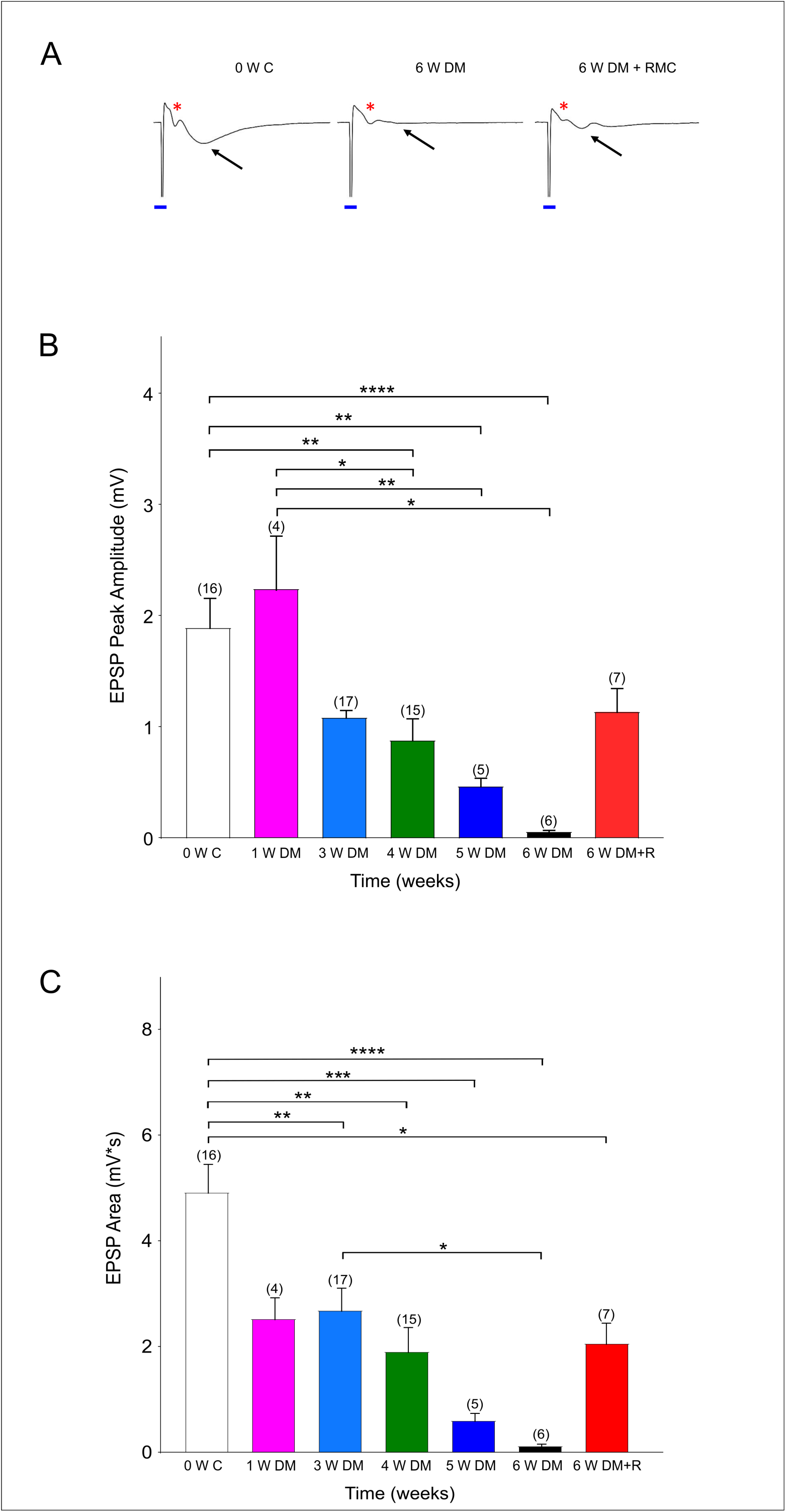
Demyelination interrupts synaptic transmission. A. Representative traces showing fEPSPs (black arrows) with a clear Afferent Volley (AV, red asterisk) at 0 Week Control (0 W C). Six week of demyelination (6 W DM) diminishes fEPSP while AV remains unchanged. Remyelination (6 W DM + RM) partly restores fEPSPs. Note the stimulus artifact that is marked with a blue line. B, C. Demyelination causes a time dependent suppression of EPSP Peak (B) and EPSP Area (C) which are partially restored with remyelination. * p< 0.05, ** p< 0.005, *** p<0.001, ****p<0.0001, One-way Anova, Bonferroni test.

### Mouse model of demyelination and remyelination

After the mice recovered from craniotomy surgery and we recorded at least one imaging session from them (7-14 days after the craniotomy surgery), we switched their diet to include 0.3% cuprizone (TD.140805, Envigo). Mice were provided access to food *ad libitum*. Food pellets were refrigerated and were replaced 3 times per week to ensure cuprizone efficiency. We continued to monitor hippocampal activity on a weekly basis. After 53 days of cuprizone diet, we identified that activity levels had reached a median level of 0 APs (Fig. 2B) in one of the mice, and we switched the mouse diet back to normal food. Mouse weight was monitored before each recording session (Supp. table 1). For electrophysiological recording experiments, mice were also injected daily with rapamycin for 12 weeks.

**Figure 2.**
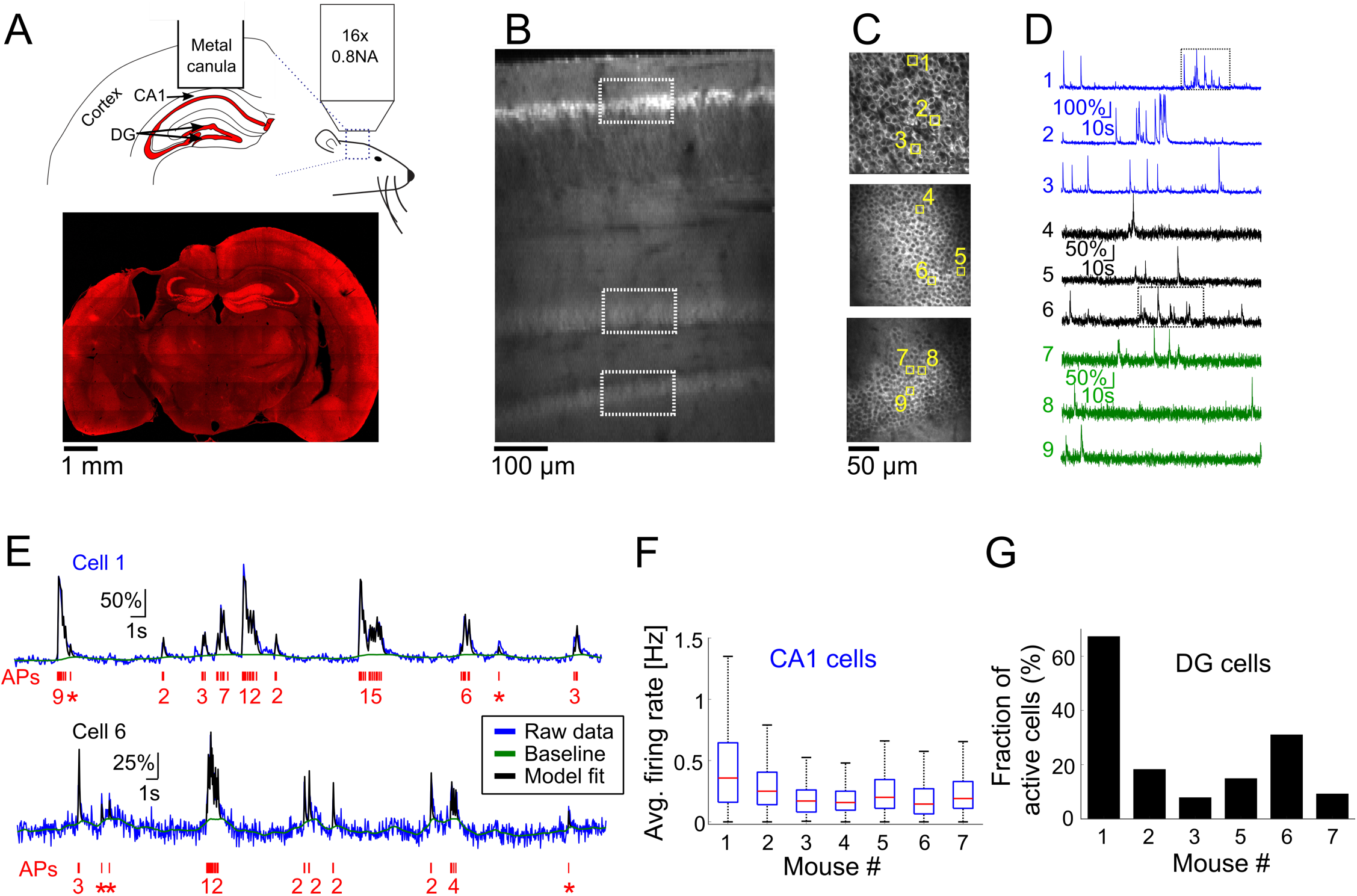
Optical recording from mouse CA1 and DG regions of the hippocampus. A. Top, schematic illustration of the mouse hippocampal recording experiments. We implanted metal cannulas with glass bottom on top of the hippocampi of transgenic Thy-jRGECO1a mice to gain optical access to CA1 and DG neurons. Hippocampus geometry was adopted from a mouse brain atlas [45]. Bottom, a coronal section from the brain of mouse 1 shows the location of the hippocampal window and the jRGECO1a expression throughout the brain. Note that some distortion to the brain tissue occurred when the metal cannula was detached from it. B. *in vivo* dorsal-to-ventral view of a Thy1-jRGECO1a line 8.31 transgenic mouse expressing the red GECI jRGECO1a in the hippocampus 3 weeks after a craniotomy surgery. The tissue on top of the hippocampal formation was removed to expose the hippocampus and a z-stack series of two-photon microscopy images shows CA1 and DG regions. White rectangles indicate the location of the CA1 and DG (upper and lower blades) projection neurons, labeled by jRGECO1a. C. Images of CA1 pyramidal cells (top image) and DG granular cells from the upper and lower blades of DG (middle and bottom images, respectively), corresponding to the respective rectangles shown in B. D. ΔF/F_0_ traces of the 9 cells highlighted with yellow squares in C. Note the differences in firing pattern and amplitude between the CA1 and DG cells. E. Model fit [27] (black line) to the raw fluorescence signal (blue line) of cells 1 and 6 (upper and lower traces, respectively, corresponding to the rectangles shown in D) to extract the baseline signal (green line) and AP firing (red, bottom, single AP events are represented with asterisks). The model was fed with jRGECO1a biophysical parameters [19], and showed good agreement to the recorded data from both CA1 (top) and DG (bottom) neurons.. F. Summary of average firing rates of all recorded CA1 pyramidal cells from 7 mice before the initiation of cuprizone diet (328-623 cells from a single recording session for each mouse, 3,167 cells total). Red lines correspond to medians, blue boxes show the 25th–75th percentile range, whisker length is the shorter of 1.5 times the 25th–75th range or the extreme data point. Outliers are not shown. G. Fraction of active DG neurons from all recorded DG neurons from 6 mice before the start of cuprizone diet (470-997 cells from a single recording session for each mouse, DG cells of mouse 4 were not visible, 3,113 cells total).

## Results

### Demyelination interrupts synaptic transmission in hippocampal CA1 neurons

Previous studies have identified that the hippocampus is severely demyelinated following 6-12 weeks of cuprizone diet, and that additional injections of rapamycin have eliminated the spontaneous remyelination that occurs when mice are fed with cuprizone [10; 28]. These mice performed poorly in a hippocampus-dependent spatial-memory task [15]. Therefore, to investigate changes in neuronal activity in the hippocampus, we conducted a set of *in vitro* experiments in acute hippocampal slices. Forty-six mice were divided into seven cohorts and control. The groups included 1, 3, 4, 5, and 6 weeks of cuprizone-diet-induced demyelination and daily rapamycin injections, and 6 weeks of cuprizone diet with rapamycin injections and additional 6 weeks of remyelination with regular diet and no rapamycin injections (Fig. 1). The control group received regular diet and daily rapamycin injections for 6 weeks, and recordings were performed at 0, 4 and 6 weeks to compare synaptic properties between demyelination and rapamycin-only conditions (Supp. Fig. 1). EPSPs together with clear AVs were evoked to monitor axon function in Schaeffer collateral commissural fibers. A successful synaptic transmission is a result of presynaptic input and a subsequent synaptic response. Sample traces shown in Figure 1 denote that in every control slice following the stimulus artefact (blue line, due to square pulse delivered with stimulation electrode), a prominent AV and EPSP were evoked (Fig. 1A, O W C, red *: AVs, black arrow for EPSPs). EPSPs presented with a peak amplitude of ~2 mV and an average area of 5 mV*s (Figure 1B, white columns, n=16). As demyelination continued, EPSP peak amplitude and area gradually started to drop becoming prominently low by 3-4 weeks (Figure 1A-B, blue and green columns, n= 17 and 15, respectively) and almost completely diminishing by 6 weeks (Figure 1A, B, black columns, n= 6). These changes were specific to demyelination because recordings in 4- and 6-weeks control groups were similar to 0 W control values (Supp. Fig. 1). Remyelination partially restored EPSPs, albeit with smaller amplitude compared to control group (Fig. 1A, B, C, 6 W DM +RM, n= 7). Therefore, demyelination effectively interrupted synaptic transmission characterized with loss of EPSPs despite sustained presynaptic input while remyelination partially restored synaptic transmission.

### Chronic optical recording of neuronal activity from CA1 and DG cells of Thy1-jRGECO1a mice

In order to further investigate the prevalence and time course of CA1 neuronal activity modulations, we conducted additional experiments for longitudinal optical recording of GECI signal from the hippocampus of lightly-anesthetized mice (n=7 mice, 10-17 weeks old when implanted with an optical window, mice 1, 4, and 7 were males; mice 2, 3, 5, and 6 were females). Optical recording from CA1 cells usually requires removal of the cortical tissue on top of the recorded area [24; 25; 29]. Recordings from the DG area also required removal of the CA1 region, causing severe damage to the hippocampal circuit [30; 31]. Recent developments in the field of three-photon functional microscopy offer deeper tissue penetration to depths greater than 1 mm, enabling the recording of CA1 cells under the intact cortex [32]; however, this method is not yet able to probe deeper structures such as the DG. To allow deep-hippocampal recording while minimizing disruption to its integrity, we followed previous studies that used red GECIs to facilitate deeper TPLSM imaging, which allows recording from the DG without removing the CA1 layer [19; 33]. We used a recently-developed transgenic line, Thy1-jRGECO1a line GP8.31, which expresses the state-of-the-art red GECI jRGECO1a under the Thy1 promoter. jRGECO1a is expressed mostly in projection neurons across various brain regions, including the CA1 pyramidal and DG granular layers of the hippocampus [21]. The enhanced imaging depth and high sensitivity achieved with jRGECO1a compared to green GECIs like GCaMP that we previously demonstrated for cortical imaging [19] enabled us to record, in 6 out of the 7 mice used in this study, from the upper and lower blades of the DG down to 650 μm under the hippocampal surface using up to 120 mW of 1100 nm excitation light (Fig. 2A-D).

Recording of spontaneous activity from CA1 and DG neurons of lightly-anesthetized mice revealed different firing patterns across the two populations (Fig. 2D, Supp. Movies 1-2). To quantify activity patterns from the recorded fluorescence traces, we used a physiological model for extracting AP firing from calcium signals (Fig. 2E) [27]. The majority of CA1 cells fired bursts of APs with a median average firing rate of 0.21 Hz and a maximal instantaneous firing rate of 8.15 Hz (3,114/3,167 CA1 cells were detected with at least firing of 1 AP; 2,099/3,167 cells fired bursts of APs, instantaneous firing rate was measured for 1 sec time bins. Data from n=7 mice, one recording session for each mouse, Fig. 2F). Most DG cells were not active during our recordings, and firing rates were substantially lower than those of CA1 cells (1,048/3,113 DG cells were detected with at least firing of 1 AP; only 184 cells from which were detected with average firing rate higher than 0.05Hz; n= 6 mice, one recording session for each mouse, Fig. 2G).

Chronic activity recording from DG neurons proved to be more challenging than CA1 recording, since the deeper location of the DG resulted in a lower signal-to-noise ratio, caused by attenuation of the incoming laser beam and outgoing fluorescence signal. Moreover, we found that in some of the mice, the recorded image quality gradually degraded 6-12 weeks after cranial window implantation (Supp. Fig. 2, the DG of mouse 4 could not be recorded, the contrast of mouse 3’s DG neurons has deteriorated one month after the craniotomy surgery, and was excluded from the rest of the study), resulting in reduced sensitivity to AP detection, or even an inability to identify DG cells. We quantified this degeneration by measuring the contrast between the cytosolic jRGECO1a signal and the nuclear signal, which should be much weaker in jRGECO1a-expressing neurons [19]. The contrast of CA1 neurons was 2-5 times higher than DG neurons in the same animal. Data from DG neurons, recorded on dates with very low contrast, were excluded from our analysis (Supp. Fig. 2B).

### Demyelination reduced CA1 projection neurons firing rate

To explore the reversible change in CA1 excitability (Fig. 1) and to quantify how demyelination and remyelination affect the activity of single neurons in the hippocampus, we fed n=4 mice with 0.3% cuprizone diet (n=3 mice for 53 days, mouse 4 optical window quality deteriorated after 16 days of cuprizone diet and it was subsequently excluded from the rest of the study, Fig. 3A). A control group of n=3 mice were implanted with a hippocampal window and received normal diet. No rapamycin was injected in order to allow for faster remyelination after the end of the cuprizone diet period [10]. Weekly monitoring of CA1 and DG activity was conducted before, during, and after the cuprizone diet period, in order to identify demyelination- and remyelination-induced changes to activity patterns in the same animal. Shortly after the initiation of the cuprizone diet, the average firing rate of CA1 cells was significantly decreased by 57% (comparing the mean of median firing rate values per mouse of n=4 mice, with 1,695 and 1,823 identified neurons on days 0 and 9 for cuprizone diet, respectively, P<10^−13^ for the decrease in each mouse firing rate, Wilcoxon Rank Sum Test; P=0.02, paired t-test between median firing rates of mice 1-4 on days 0 and 9; Fig. 3B-C, Supp. table 1). During the 53 days of cuprizone diet, CA1 median firing rate significantly decreased for individual mice (P=0.002, 0.064, 0.035, Mann-Kendall Trend Test for mice 1, 2, and 3, respectively, Supp. Fig. 3), but the group effect was not significant, presumably because of the small sample size (P=0.07, paired t-test for change for median firing rates of mice 1-3). We also detected short-term increases in firing rates (Fig. 3B-C, days 20-30) that might be associated with partial spontaneous remyelination, as was previously identified in cuprizone-treated mice [10]. On the last day of the cuprizone diet, the average firing rate was significantly reduced to 16.5% of pre-diet levels (comparing the mean of median firing rate values per mouse of n=3 mice, with 1,135 and 1,356 identified neurons on days 0 and 53 for cuprizone diet, respectively. P<10^−36^, Wilcoxon Rank Sum Test, Supp. Movies 1,3). A similar decrease was identified in the fraction of cells that fired bursts on APs (Fig. 3D, Methods). Monitoring CA1 activity of n=3 control mice showed no similar decrease. Changes in firing rates and burst activity were smaller than in the cuprizone group, had no identified monotonous trend, and presumably were the result of differences in anesthesia levels and brain state (Fig. 3E-G, Supp. Movies 4,5; P=0.99, 0.86, 0.14, Mann-Kendall Trend Test for mice 5, 6, and 7, respectively; P=0.53 and 0.49, paired t-test for change of median firing rates of mice 5-7 between days 0 and 7, and days 0 and 49, respectively; Supp. Fig. 4). Repeated ANOVA test between the cuprizone and control groups for comparing effects of time and group yielded no statistically significant difference in the group and the interaction between the time and group, respectively results (P=0.03 for the time effect, p=0.77 for the group effect, and P=0.18 for the group-time interaction effect).

**Figure 3.**
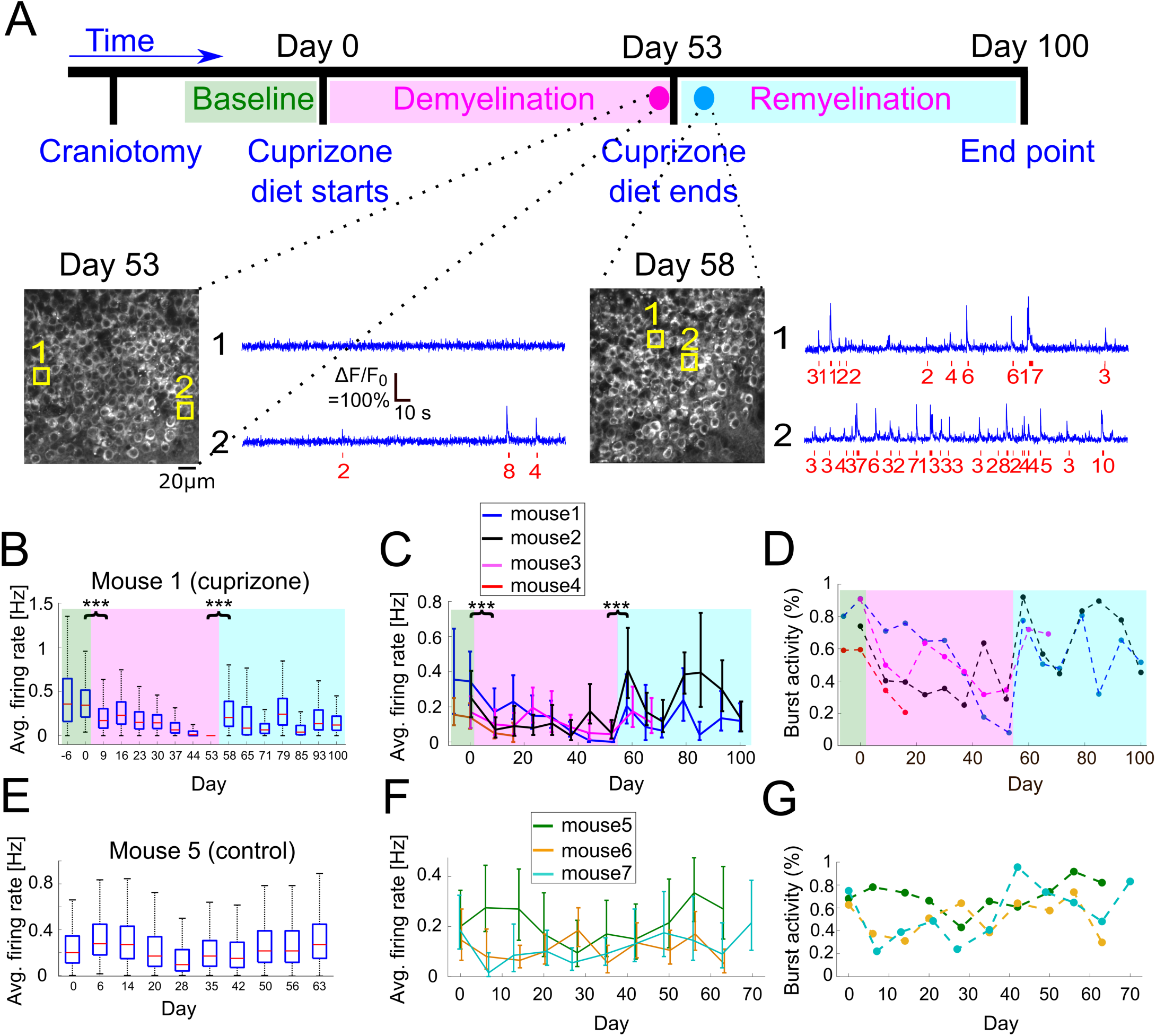
Changes to CA1 activity patterns during demyelination and remyelination. A. Time line of the demyelination/remyelination experiment for mice 1-4. Craniotomy surgery was performed between day −14 to −7, and baseline activity recording from CA1 and DG neurons started between day −6 to day 0 (indicated by green background in panels B-D). On day 0, after an activity recording session, we switched the diet to include 0.3% cuprizone. This diet was administered until day 53 (demyelination period, indicated by magenta background in panels B-D), while we maintained weekly activity recordings. On day 53, after an activity recording was completed, we switched the mice back to normal diet and monitored activity until day 100 (remyelination period, indicated by cyan background in panels B-D). Bottom, two example fields of view acquired from mouse 1 (cuprizone group), showing the CA1 cells and activity of two example cells on days 53 and 58. The estimated timing and number of APs is shown (red) beneath each raw data trace (blue). B. Summary of the average firing rates from all recorded CA1 neurons from mouse 1 during 107 days of recording. Firing rates decreased significantly from 0.345 Hz to 0.17 Hz between day 0 and day 9 (P=1.8*10^−37^, Wilcoxon Rank Sum Test), after cuprizone diet had been introduced. Activity levels continued to significantly decrease down to zero on day 53 (P=0.002, Mann-Kendall Trend Test), and were significantly increased to 0.205 Hz on day 58, 5 days after normal diet was resumed (P=4*10^−133^, Wilcoxon Rank Sum Test). Activity levels on the following days continued to increase compared to the demyelination period (P=0.064, Mann-Kendall Trend Test), but remained lower than the pre-cuprizone period. Magenta and cyan backgrounds highlight the demyelination and remyelination periods, respectively; red lines correspond to medians, blue boxes show the 25th–75th percentile range, whisker length is the shorter of 1.5 times the 25th–75th range or the extreme data point. Outliers are not shown. *** - P<0.001. C. Summary of the average firing rates from all recorded CA1 neurons from all cuprizone-treated mice. Solid lines connect the distribution medians, and the error bars indicate the 25^th^ to 75^th^ percentile range for each recording session. For all recorded mice, there was a significant decrease in the average firing rate upon the start of cuprizone diet, between day 0 and day 9 (n=4 mice, P<10^−13^, Wilcoxon Rank Sum Test), and a significant increase upon the termination of cuprizone diet, between day 53 and day 58 (n=3 mice, P<10^−32^, Wilcoxon Rank Sum Test). Note that the rapid increase in firing rate on day 58 was followed by a second decrease in activity rate. *** - P<0.001. D. Summary of the fraction of cells that fired bursts of APs from all recorded cells for all cuprizone-treated mice. This fraction decreased for all mice during the demyelination period and rapidly recovered upon the start of the remyelination period. Colors are the same as in panel C. P-values for the decrease were 0.035, 0.108, and 0.064 for mice 1, 2, 3, respectively (Mann-Kendall Trend Test). E. Summary of the average firing rates from all recorded CA1 neurons from control mouse 5, which received normal diet throughout the recording period, shows no similar decreases or increases in firing rate as those recorded from cuprizone-treated mice. F. Summary of the average firing rates from all recorded CA1 neurons from all control mice. Solid lines connect the distribution medians, and the error bars indicate the 25^th^ to 75^th^ percentile range for each recording session. None of the traces showed significant monotonous trend (P=0.99, 0.86, 0.14, for mice 5, 6, 7, respectively, Mann-Kendall Trend Test). G. Summary of the fraction of cells that fired bursts of APs from all CA1-recorded neurons from all control mice. Colors are the same as in panel F.

**Figure 4.**
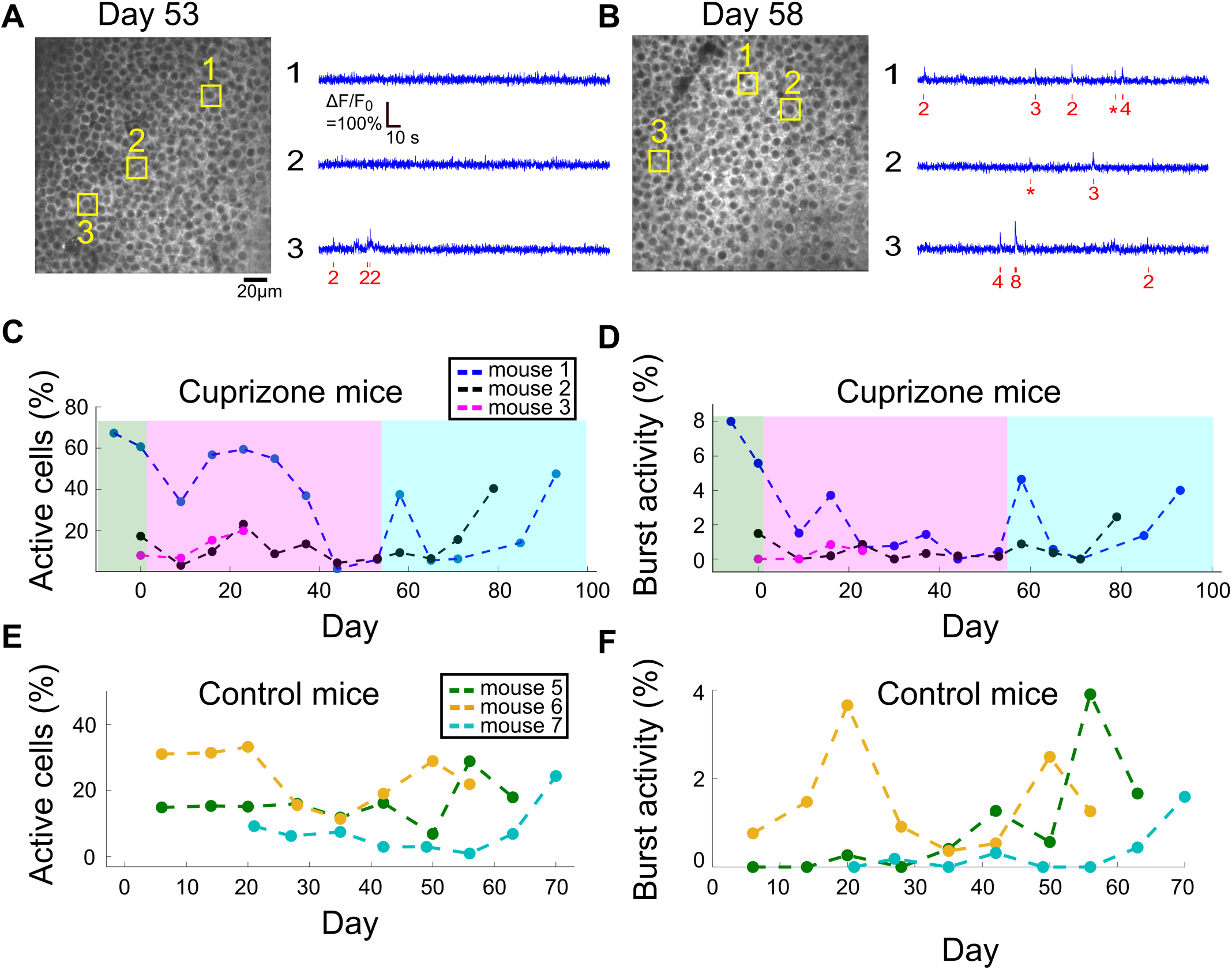
Changes to DG activity patterns during demyelination and remyelination. A. An example field of view of DG neurons from mouse 1, after 53 days of cuprizone diet, with activity traces of three example cells. The estimated timing and number of APs is shown (red) beneath each raw data trace (blue). B. An example field of view of DG neurons from mouse 1, 5 days after the end of the cuprizone diet, with activity traces of three example cells. The estimated timing and number of APs is shown (red) beneath each raw data trace (blue). C. Summary of the fraction of active DG cells from all recorded DG cells for all cuprizone-treated mice. The low-level recorded activity of DG cells is in line with previous findings [33]. However, the cuprizone diet reduced the activity even further, and the termination of cuprizone diet was followed by a noticeable recovery. DG cells of mouse 3 could be recorded only during the first 4 recording sessions. D. Summary of the fraction of DG cells that fired bursts of APs from all recorded DG cells in cuprizone-treated mice shows a similar trend to the change in CA1 activity patterns. Colors are the same as in panel A. E. Summary of the fraction of active DG cells from all recorded DG cells for all control mice. F. Summary of the fraction of DG cells that fired bursts of APs from all recorded DG cells in control mice. Colors are the same as in panel C.

### Spontaneous remyelination restored the activity of CA1 excitatory neurons

It has been shown multiple times that once cuprizone diet is stopped and mice are returned to their normal diet, spontaneous remyelination begins [8; 10; 11; 15]. Therefore, once we stopped the cuprizone diet and returned the mice to normal diet, we kept monitoring the same mice during the remyelination period and identified a substantial and significant increase in the average firing rate of individual mice within 5 days after the end of the cuprizone diet, but the group effect was not significant (1,356 and 1,207 neurons from n=3 mice recorded on days 53 and 58, respectively. P<10^−32^ for the change in individual mice firing rate, Wilcoxon Rank Sum Test; P=0.08, paired t-test for change in median firing rates; Supp. Movies 3,6, Supp. Fig. 3), which was animal-dependent in magnitude, but returned overall activity back to similar levels as the pre-cuprizone period (−40.5%, +4.8%, and +104% of the pre-cuprizone median firing rate, n=3 mice, 1,135 and 1,207 neurons from days 0 and 58, respectively). Interestingly, this recovery wasn’t monotonous (non-significant results in Mann-Kendall Trend Test for days 53-100), and included a second cycle of decreasing and increasing of CA1 cell activity (Fig. 3B-C, days 65-70). We monitored cellular activity for 47 additional days following the termination of cuprizone diet, and identified partial recovery of cellular firing rates to lower levels than before the start of cuprizone diet (35% and 52% of the activity level on day 0, data from 807 and 1,201 neurons from n=2 mice, measured on day 0 and day 100, respectively; data shown in Fig. 3C-D, Supp. Movie 7, Supp. Fig. 4).

### Demyelination reduced activity levels of DG excitatory neurons

For all tested mice, DG neuronal activity levels were lower than those of CA1 cells, and in the majority of DG neurons, firing of APs was not detected under our experimental conditions, similar to previously-reported data [33]. In neurons that AP firing was detected, most fired a low number of APs. Therefore, we also quantified DG activity levels by assessing the fraction of cells that fired any number of APs, and the fraction of cells that fired at least one burst of more than 5 APs within a 660 ms time bin. Cuprizone diet caused a rapid decrease in the fraction of active cells by 49%, and an 89% decrease in bursting activity, but insufficient group size resulted in non-significant p-values (1,977 and 2,326 neurons recorded on days 0 and 9 respectively, n=3 mice; P=0.19 and 0.26, t-test for change in fraction of active and bursting cells, respectively). Average firing rate was significantly decreased for 2 out of 3 mice (P<10^−15^ for mice 1 and 2, P=0.338 for mouse 3, Wilcoxon Rank Sum Test). This decrease in activity levels continued during the cuprizone diet period, similar to the trend detected in CA1 cells (Fig. 4A-B, Supp. Fig. 5). On the last day of the cuprizone diet, the fraction of active cells was decreased to 15% of its value on day 0 (1,419 and 1,334 neurons from n=2 mice, recorded on day 0 and 53, respectively, P<10^−10^ for change in average firing rate, Wilcoxon Rank Sum Test, Supp. Movies 2,8). Activity recording in control mice did not show such a decrease. (P=0.59 and 0.30, paired t-test for change in fraction of active and bursting cells between first and second recording dates, respectively; Fig. 4C-D, Supp. Movies 9,10, Supp. Fig. 6).

### Spontaneous remyelination restored the activity of DG projection neurons

DG neurons presented a similar recovery pattern as CA1 neurons. DG cell activity increased rapidly upon termination of cuprizone diet, from 15% to 59% of the fraction of active cells on day 0 (1,419, 1,334, and 1,059 cells from n=2 mice on days 0, 53, and 58, respectively, Supp. Movie 11, Supp. Fig. 5). This was followed by a second decrease and increase in activity levels, to comparable levels to day 0 (78.3% and 220.2% of fraction of active cells on day 0 for mice 1 and 3, respectively, data from 1,419 and 1,657 cells on day 0 and on the last day of recording, respectively; Fig. 4A-B, Supp. Movie 12, Supp. Fig. 6).

## Discussion

Repeating cycles of demyelination and remyelination are associated with the progression of MS in the majority of patients, who experience the relapsing-remitting form of the disease [34]. This highlights the necessity to understand their impact upon brain activity. In this study, we measured the effects of cuprizone-induced demyelination and remyelination on hippocampal activity, which recapitulates the loss and partial recovery of myelin, consistently in the same brain regions, and with a reliable and reproducible time course [10; 15; 28]. We combined electrophysiology and functional microscopy for acute and long-term monitoring of the activity of projection neurons in CA1 and DG regions with cellular resolution over more than 100 days, during both the demyelination and remyelination phases. We identified cuprizone-induced deterioration of synaptic transmission of CA1 cells, and quantified the longitudinal effect of this deterioration on hippocampal activity *in vivo*. We report large changes in neuronal firing rates, with both fast and long-term components that followed the initiation and cessation of cuprizone diet. These changes were associated with the demyelination and remyelination processes, as were previously reported, and are in agreement with behavioral deficits and recovery that were also reported in cuprizone mice [10; 28], and changes to the excitability level of single neurons [35; 36], as well as disrupted large-scale activity patterns [37].

The time-scales of cuprizone-mediated brain activity degeneration, and recovery once cuprizone diet was stopped, are in agreement with previous studies [10; 28]. However, our recordings have identified rapid changes and non-monotonous trends for both conditions that have not yet been reported. A large fraction of the activity change occurred within 5-9 days (one recording session) from the initiation and cessation of cuprizone diet, faster than the reported time scale of 4-6 weeks for demyelination and remyelination [10]. Interestingly, this relatively rapid change was followed by a “rebound” change in activity levels, i.e. the activity increased during cuprizone diet, and decreased during remyelination. Increased activity during demyelination, approximately 3-6 weeks after the start of cuprizone diet, may be explained by partial remyelination due to maturation of the oligodendrocyte progenitor cells into mature myelinating oligodendrocytes [10], but other characteristics of the effects of cuprizone on neuronal and non-neuronal cells, and how it translates to the reported rapid changes in activity, will require additional studies. It is possible that cuprizone has, in addition to the reported effect on oligodendrocytes, a direct effect on neuronal health, or an indirect effect, mediated by glial cells, that resulted in the rapid changes we identified. In addition, this study included both male and female mice, and the reduction in activity level seems to be stronger in males (mice 1 and 4). Though the number of mice used in this study do not allow for a definitive assessment of sex-specific differences, such differences may be expected based on the existing findings [8; 38].

In agreement with fluorescent imaging, electrophysiological recording of CA1 synaptic transmission showed significant depression staring by 3 weeks of demyelination, earlier than previously reported [10]. Depression of EPSPs progressed on a weekly basis reaching a nadir level by six weeks of demyelination. Interestingly, afferent volley was well preserved even in 6-week demyelinated slices suggesting that the primary site of failure for synaptic transmission is due to post-synaptic elements. Indeed, cuprizone-induce demyelination reduces AMPA receptors in the CA1 region of hippocampus [15]. Additionally, deficits in excitatory synaptic transmission during experimental autoimmune encephalomyelitis (EAE) correlated with disruption of PSD-95 integrity involving both AMPA and NMDA receptor-mediated currents [39; 40] while preserving presynaptic function in CA1 neurons. In contrast, recordings in layer 5 pyramidal neurons in somatosensory cortex of mice that were kept on cuprizone diet for 5 weeks revealed that action potential (AP) propagation switched from rapid saltatory towards a slow continuous wave broadening the presynaptic AP with reduced velocity presumably due to redistribution of fast acting K channels [36]. Furthermore, AP changes were accompanied with demyelination of internodes, variable reorganization of nodal domains, and frequent sprouting of axons. Taken together, these findings suggest that loss of myelin in gray matter causes a wide range of site-specific structural and functional changes.

Optical activity monitoring allows probing how connected brain regions are affected by external input, and how they affect each other. We recorded activity both from CA1 and DG neurons in the same mice, and identified substantial differences in their activity patterns (Figs. 2-4), in agreement with previous studies [29; 33]. The deep-tissue location and low level of activity of DG neurons under anesthesia make them more challenging to monitor compared to CA1 neurons (Supp. Fig. 2). Therefore, more studies are required for better understanding cuprizone-induced changes in DG cells, preferably in awake mice, where activity levels are expected to be higher [33]. Nevertheless, longitudinal DG recording is feasible using laser power that is not expected to produce substantial heating or thermal damage to the tissue [41]. Our electrophysiological studies have highlighted that cellular excitability of CA1 neurons was interrupted during cuprizone-induced demyelination, suggesting impaired synaptic transmission which was restored during remyelination. Future studies may characterize how DG pre- and post-synaptic activity is modulated and how it affects the CA1 pre-synaptic signal via the feed-forward hippocampal circuitry. Such studies may reveal whether cuprizone-mediated diet attenuates activity in other brain regions, and hence also the input to the hippocampus, reduces the excitability of DG neurons via changes in excitation-inhibition balance, or similarly affects the cellular excitability in both CA1 and DG neurons.

A limitation of the *in vivo* approach we used in this study is the relatively low number of mice in each group, which is related to the amount of time required to record and analyze the activity data from each mouse. The resulted small group size limited the significance of the statistical analysis we performed. We received significant p-values for changes of activity characteristics from the same animal, which relied on the large number of cells we sampled. However, our statistical analysis was less conclusive for t-tests across median values from mice in the same group, and non-significant for repeated ANOVA test across the cuprizone and control groups. A preferred approach for overcoming this limitation is to increase the group sizes, which might require adopting an automatic analysis pipeline for improving the throughput of such experiments [42].

Finally, applying optical methods for monitoring the brain condition offers an attractive approach for acquiring essential *in vivo* information to link structure, function, and behavior. We demonstrated that TPLSM recording of GECI signal enables direct evaluation of functional properties of neurons in a mouse model for MS-like symptoms. Moreover, we show that deterioration of synaptic transmission in CA1 cells is reflected in the activity levels of these neurons, and can be measured *in vivo* in a longitudinal manner. Future studies may combine this approach with emerging methods for *in vivo* measurements of the myelin sheath condition using label-free third harmonic generation [43; 44] to allow for all-optical detection of demyelination and activity levels in the same cells. Complementary behavioral data can be acquired with standard behavioral and cognitive tests for assessing the mouse condition, such as the Morris water maze [15]. Such experimental platforms will provide a more holistic assessment of the mouse condition. Such experimental platform may also be used as an evaluation testbed for comparing the efficacy of new pharmaceutical intervention treatments on protecting synaptic transmission and neuronal activity patterns, reducing demyelination levels, and relieving behavioral deficits.

## Acknowledgements

We thank the HHMI Janelia GENIE project for developing and sharing the Thy1-jRGECO1a transgenic mice. We thank Dr. Vanzetta and his colleagues for developing and sharing the AP extraction model used in this study. The authors would like to thank Drs. Albert Lee and Jae Sung Lee for sharing their hippocampal window implantation technique, Drs. Ranjan Dutta, Chris Nelson, and Dimitrios Davalos for reviewing the paper and for their insightful comments, and Sarah Stanko and Anthony Chomyk for assisting with mouse husbandry. This research was partially funded by an MS Society Pilot Grant (PP-1901-33093) to HD and NIH grant (R35NS09730) to BT. Materials and data for SB’s part were provided by the Cleveland Clinic Foundation (CCF). All rights, title, and interest in the materials and Data are owned by the CCF.

