## Supplementary Figures for "Reversible Loss of Hippocampal Function in a Mouse Model of Demyelination/Remyelination"

**Supplementary figure legends**

**Supplementary Figure 1**

Time-dependent demyelination-mediated changes in synaptic transmission.

(A, B) Bar plots denote that the synaptic transmission is due to cuprizone mediated demyelination while synaptic properties of control group (receiving daily Rapamycin injections) remains the same. * $p < 0.05$, ** $p < 0.005$, *** $p<0.001$, ****$p<0.0001$, One-way Anova, Bonferroni test.

**Supplementary Figure 2**

A. Example images of three field of views acquired from CA1 (upper row) and DG (lower row) layers of the same mouse (mouse 2) over the course of 100 days.

B. The contrast of individual neurons was calculated as the fractional difference between their somatic and nuclear signals (Methods), as illustrated in the upper-left image. The median contrast across all recorded cells from the same mouse is shown for CA1 and DG cells for all recording sessions. Green circles indicate recording sessions with low DG cells contrast, that were excluded from the analysis.

**Supplementary Figure 3**

Example fields of view of CA1 neurons, acquired from mouse 1 on days 0 (before starting cuprizone diet), 53 (last day of cuprizone diet), 58 and 100. For each field of view, activity traces of 5 example cells are shown (blue traces) with the estimated timing and number of APs (red ticks and numbers, single AP events are represented with asterisks). The cell locations in each field of view are indicated by yellow rectangles.

**Supplementary Figure 4**

Example fields of view of CA1 neurons, acquired from mouse 5 (control group) on days 0, 51, 56, and 63. For each field of view, activity traces of 5 example cells are shown (blue traces) with the estimated timing and number of APs (red ticks and numbers, single AP events are represented with asterisks). The cell locations in each field of view are indicated by yellow rectangles.

**Supplementary Figure 5**

Example fields of view of DG neurons, acquired from mouse 1 on days 0 (before starting cuprizone diet), 53 (last day of cuprizone diet), 58 and 93. For each field of view, activity traces of 5 example cells are shown (blue traces) with the estimated timing and number of APs (red ticks and numbers, single AP events are represented with asterisks). The cell locations in each field of view are indicated by yellow rectangles.

**Supplementary Figure 6**

Example fields of view of DG neurons, acquired from mouse 5 (control group) on days 0, 51, 56, and 63. For each field of view, activity traces of 5 example cells are shown (blue traces) with the estimated timing and number of APs (red ticks and numbers, single AP events are represented with asterisks). The cell locations in each field of view are indicated by yellow rectangles.

### Supplementary Figure 1

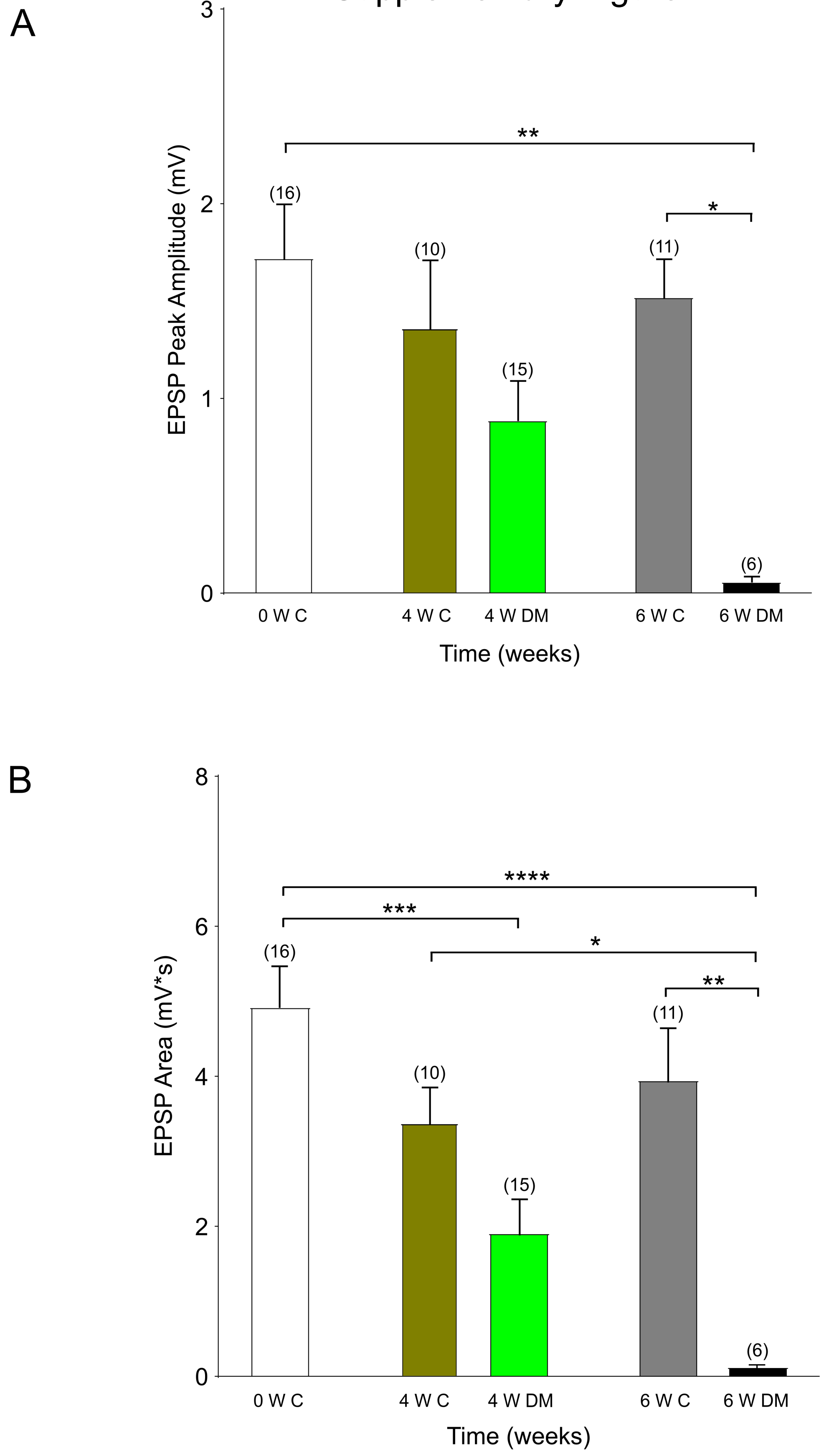

### Supplementary Figure 2

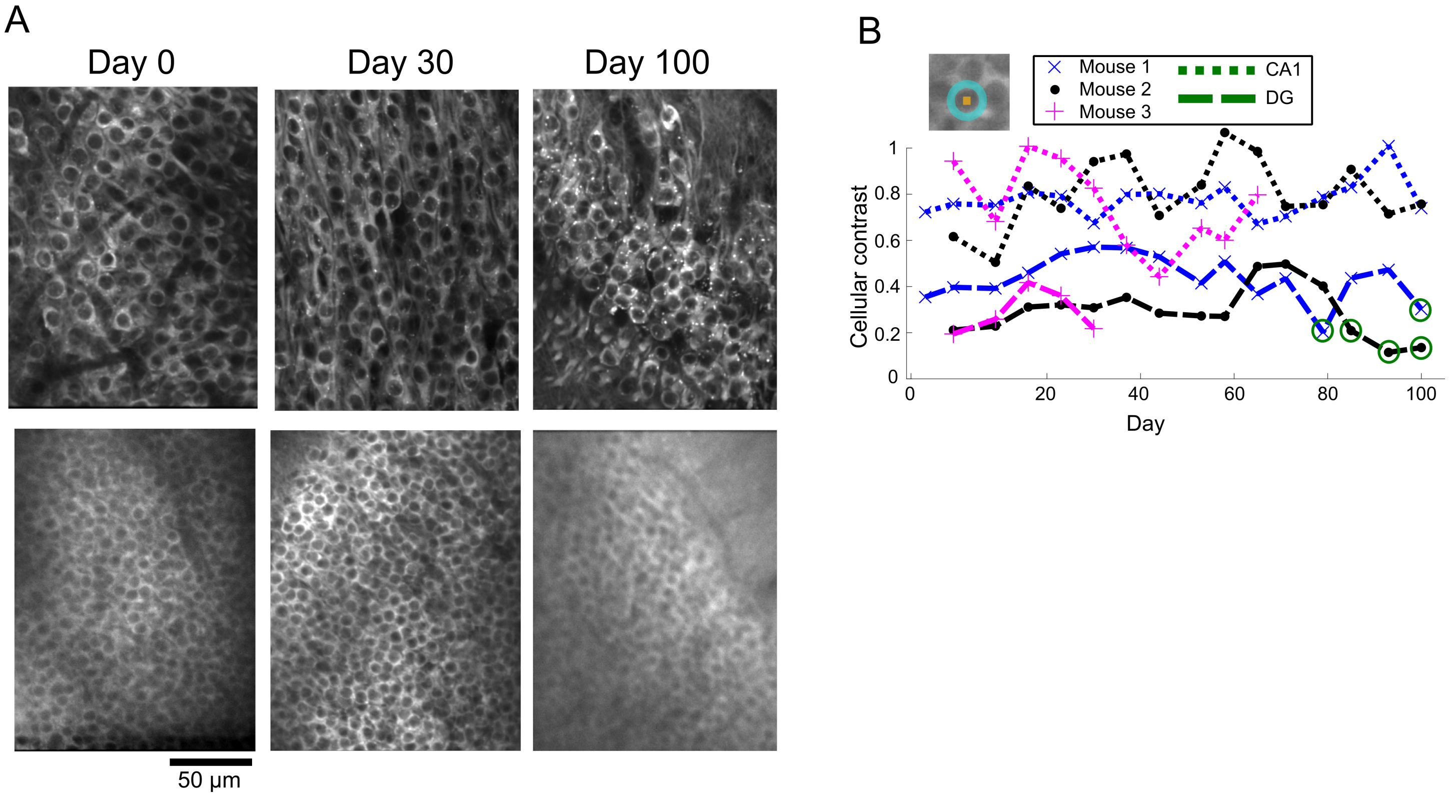

### Supplementary Figure 3

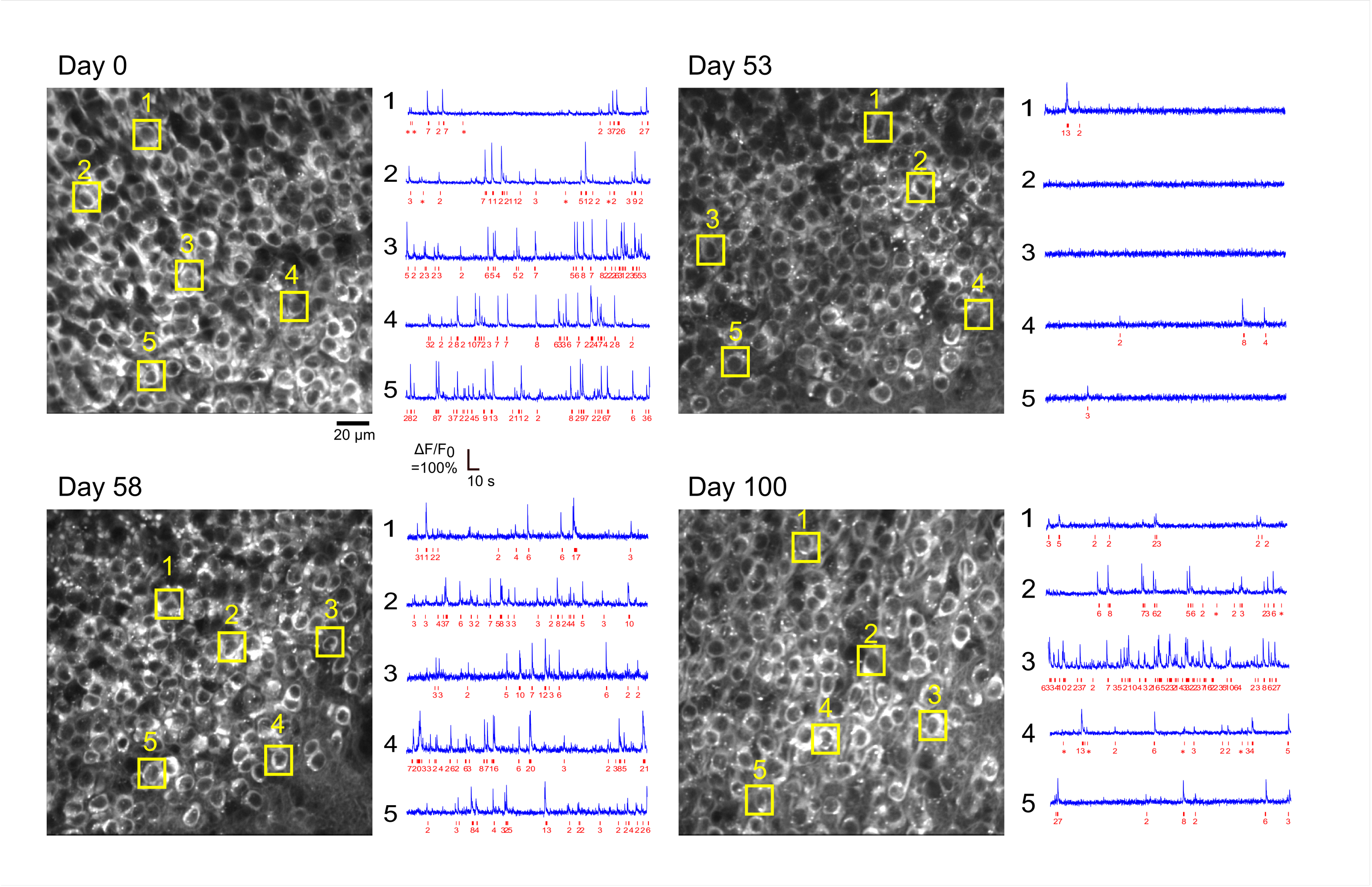

### Supplementary Figure 4

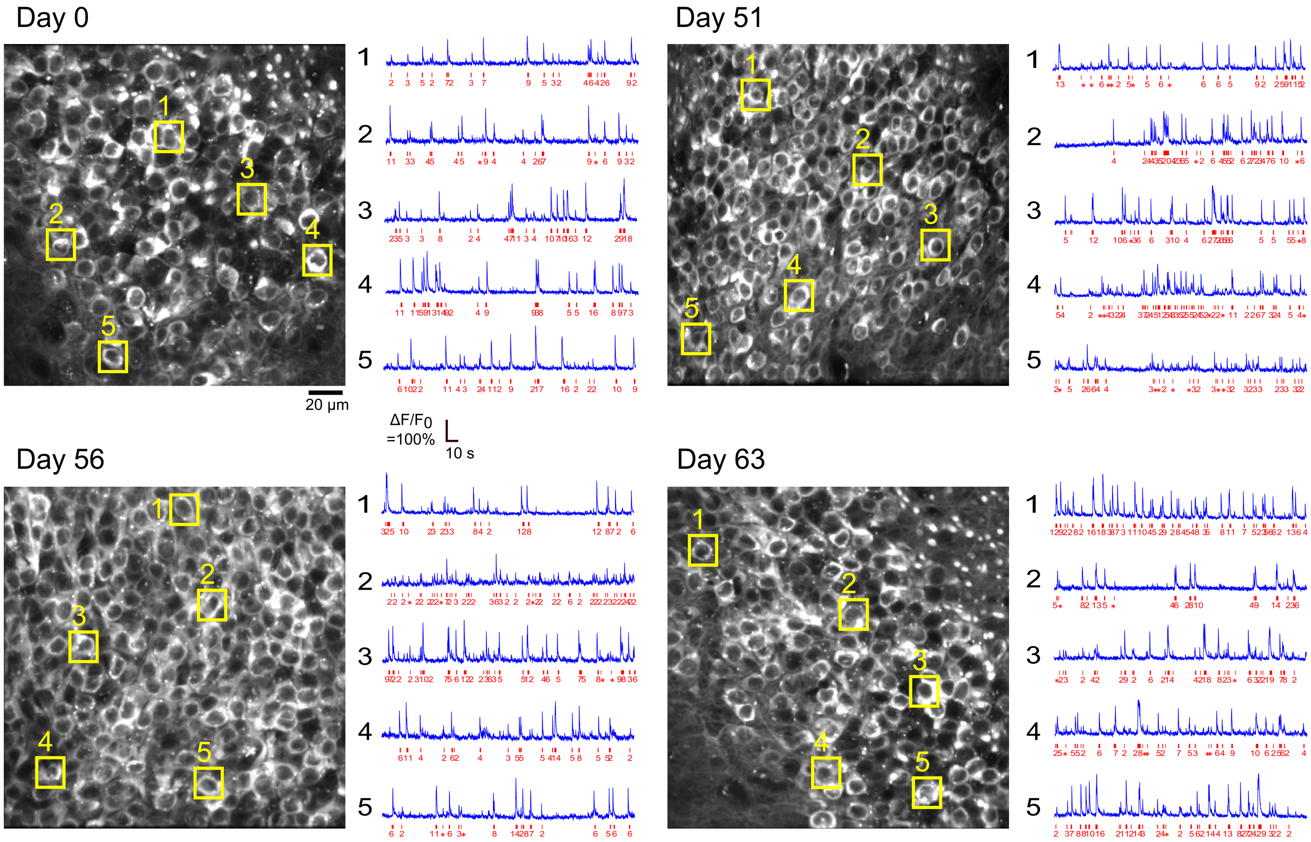

### Supplementary Figure 5

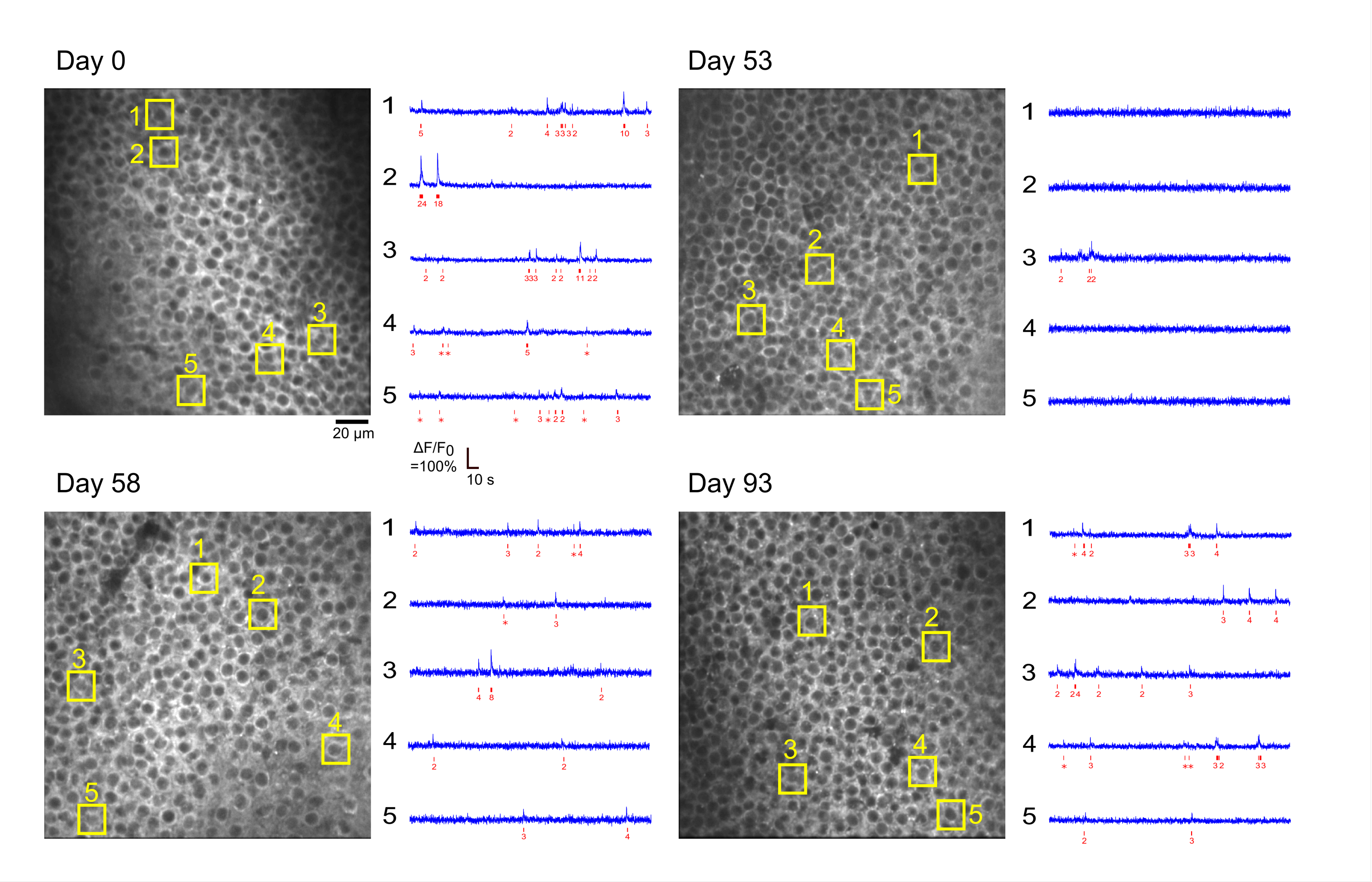

### Supplementary Figure 6

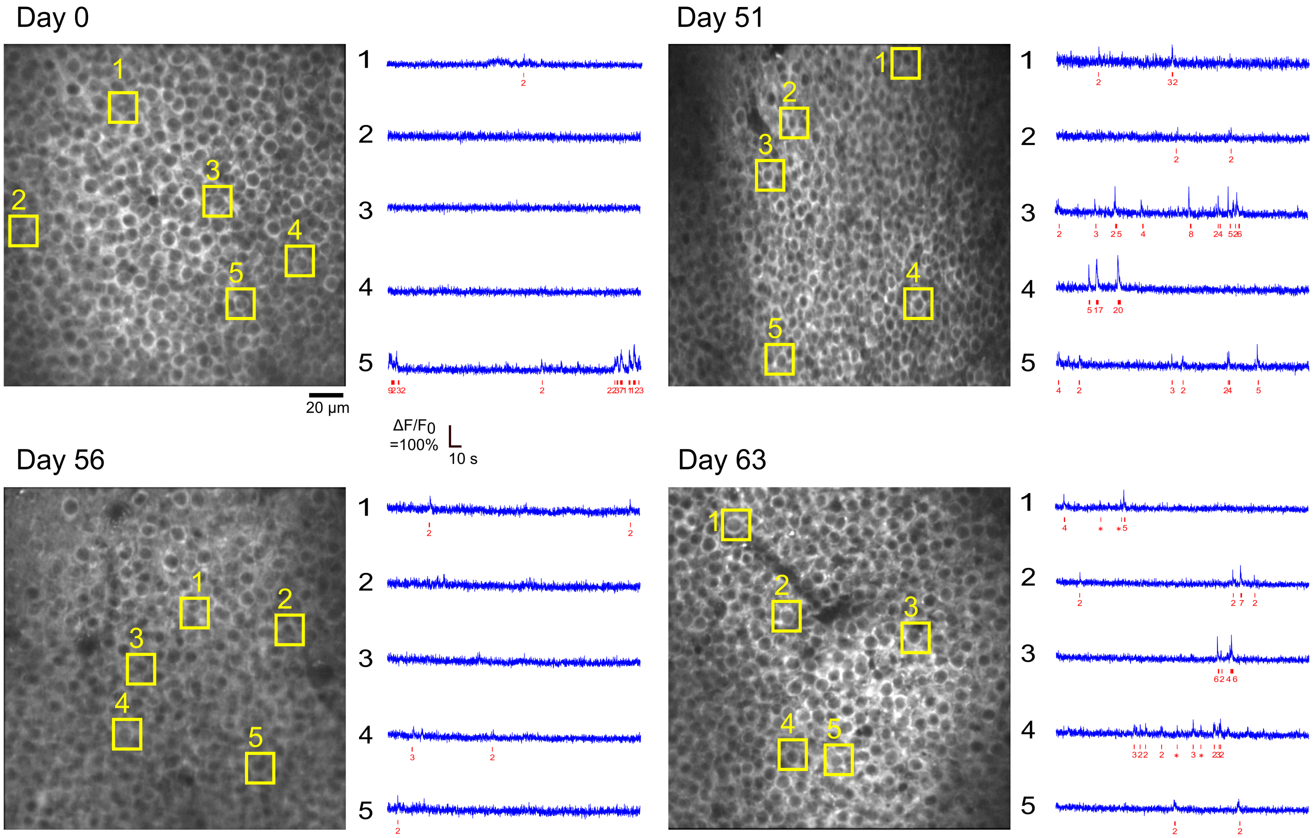
